# PAMG-AT: A Physiological Attention Multi-Graph Model with Adaptive Topology for Stress Detection using Wearable Devices

**DOI:** 10.64898/2026.03.02.709179

**Authors:** Osman Yildiz, Abdulhamit Subasi

## Abstract

Stress detection with wearable physiological sensors is vital in digital health and affective computing. Conventional machine learning techniques usually examine physiological signals separately, missing the intricate inter-signal connections involved in the human stress response. While deep neural networks offer high accuracy, they function as black boxes, offering minimal understanding of the physiological processes behind stress detection. This study introduces a hierarchical graph neural network framework for WESAD stress detection, establishing a methodology for affective computing that emphasizes interpretability and extensibility while maintaining strong predictive performance. We proposed PAMG-AT (Physiological Attention Multi-Graph with Adaptive Topology) which is a hierarchical graph neural network architecture, for stress detection using multimodal physiological signals. In this framework, physiological features serve as nodes within a knowledge-driven graph, while edges represent established physiological relationships, including cardiac-electrodermal coupling and cardio-respiratory interaction. The architecture employs a three-level attention mechanism: spatial encoding via Graph Attention Networks (GAT) to assess feature importance, temporal modeling with a Transformer to capture dynamics across time windows, and global pooling for classification. The model is evaluated using three sensor configurations (chest-only, wrist-only, and hybrid) on the WESAD dataset, employing rigorous Leave-One-Subject-Out (LOSO) cross-validation. PAMG-AT achieves competitive performance, with 94.59% accuracy (±6.8%) for chest sensors, 91.76% (±9.2%) for wrist sensors, and 92.80% (±8.33%) for the hybrid configuration. The proposed method provides interpretability via attention weights, revealing that ECG-EDA relationships (cardiac-electrodermal coupling) are most predictive of stress. Three low-responder subjects (S2, S3, S9) with atypical physiological stress patterns demonstrate lower accuracy (81-87%), offering clinically valuable insights for personalized stress management. The effective wrist-only configuration, achieving 91.76% accuracy, supports practical deployment in consumer wearables.

## 1. Introduction

Stress-related disorders are an increasing public health issue, as chronic stress is linked to cardiovascular problems, metabolic issues, and mental health conditions (Hassard et al., 2018). Conventional methods like self-report questionnaires and clinical interviews are retrospective and lack the capability for real-time monitoring (Jui et al., 2026). Wearable sensors, which continuously track physiological responses, provide an objective way to measure stress autonomic responses, supporting automated detection systems that connect laboratory research with clinical practice (AlMakinah et al., 2025).

Physiological stress responses involve activation of the sympathetic nervous system, leading to observable changes such as a faster heart rate, lower heart rate variability, increased skin conductance, quicker and shallower breathing, and peripheral temperature shifts (Giannakakis et al., 2022; Healey & Picard, 2005). Using multiple sensors together allows for a more complete detection of this varied stress response than relying on a single type of signal (Yaseen & Şahin, 2026).

The WESAD dataset (Schmidt et al., 2018) offers a benchmark for wearable stress detection, featuring synchronized physiological data from chest-worn (RespiBAN) and wrist-worn (Empatica E4) sensors recorded during the Trier Social Stress Test. Previous studies have achieved high classification accuracy using handcrafted features with traditional machine learning methods (93.12% with LDA; Schmidt et al., 2018) and deep neural networks (95.21%; Bobade & Vani, 2020). Nonetheless, deep learning models often act as black boxes, providing limited understanding of the physiological processes behind their predictions—an important drawback for clinical use, where interpretability is vital (Rudin, 2019).

Graph Neural Networks (GNNs) provide a useful approach for analyzing physiological signals by explicitly modeling relationships between features. In particular, Graph Attention Networks (GATs) allow for interpretable results through attention weights, which identify the relationships between signals that influence classification decisions (Veličković et al., 2017). Recent studies in EEG classification (Klepl et al., 2024) and brain connectivity analysis (Ali et al., 2025) showcase the effectiveness of GNNs for biosignal analysis. Nevertheless, the application of GNNs in multimodal wearable stress detection remains largely underexplored.

A major challenge involves creating graph representations that are physiologically meaningful. The human stress response includes well-understood mechanisms such as cardiac-electrodermal coupling, cardio-respiratory interactions through respiratory sinus arrhythmia, and thermoregulatory processes, all of which can be represented as graph edges. This approach, guided by existing knowledge, works alongside data-driven methods to improve interpretability and generalization. Moreover, stress responses demonstrate both spatial relationships (connections between signals) and temporal changes (how these relationships evolve over time), requiring models that can capture both aspects.

Inter-individual variability presents an additional challenge: certain people show reduced physiological responses to stress even when experiencing psychological distress (Carroll et al., 2017). Recognizing these low-responder phenotypes has important clinical implications for personalized stress management, but previous WESAD studies mainly concentrated on overall performance metrics.

This study presents PAMG-AT (Physiological Attention Multi-Graph with Adaptive Topology), a hierarchical graph neural network for multimodal wearable stress detection. The key contributions are:

i. **Knowledge-driven graph construction**: Physiological features serve as nodes with edges representing established physiological relationships (cardiac-electrodermal coupling, cardio-respiratory interaction, thermoregulation), enabling domain knowledge integration while learning predictive connections through attention.
ii. **Hierarchical attention architecture**: Graph Attention Networks capture spatial inter-signal relationships, while Transformer encoders model temporal dynamics across time windows, jointly capturing multiscale stress response patterns.
iii. **Cross-modality fusion strategy**: For hybrid sensor configurations, explicit cross-modality edges connect corresponding signals from chest and wrist sensors, enabling principled multi-sensor integration.
iv. **Comprehensive evaluation**: Leave-One-Subject-Out cross-validation across chest-only, wrist-only, and hybrid configurations ensures generalization to unseen subjects.
v. **Interpretability analysis**: Attention weight analysis reveals which physiological relationships are most predictive, with identification of low-responder subjects providing clinically actionable insights.

The remainder of this paper is organized as follows: Section 2 reviews related work; Section 3 details materials and methods; Section 4 presents results; Section 5 discusses implications and limitations; Section 6 concludes with future directions

## 2. Literature Review

Schmidt et al. (2018) introduced the WESAD dataset, establishing baseline performance of 93.12% accuracy using Linear Discriminant Analysis with handcrafted statistical and frequency-domain features. Subsequent studies explored various classical algorithms: (Nkurikiyeyezu et al., 2019) demonstrated a significant performance gap between person-specific (98.9%) and generic (83.9%) Random Forest models, highlighting inter-individual variability challenges. (Hsieh et al., 2019) identified electrodermal activity features as particularly discriminative when combined with XGBoost gradient boosting.

Deep learning methods have advanced WESAD classification by enabling automatic feature extraction. (Malviya et al., 2023) achieved 98% accuracy using LSTM networks that capture temporal dependencies in physiological signals. (Singh et al., 2024) proposed a hybrid CNN-LSTM architecture achieving 90.45% accuracy for IoMT applications. (Ali et al., 2025) developed a CNN-BiLSTM model for PPG-based stress monitoring, achieving 94.1% accuracy with wrist-worn sensors.

Attention mechanisms have emerged as powerful tools for stress detection. (Tanwar et al., 2024) integrated attention with CNN-LSTM architectures, achieving 92.7% accuracy through dynamic prioritization of temporal features. Transformer-based approaches have shown particular promise: (Wu et al., 2023)applied self-supervised multimodal Transformers to EDA, BVP, and temperature signals, while (Oliver & Dakshit, 2025) achieved 99.73-99.95% accuracy using single-modality Transformers with subject-specific validation. (Goetz et al., 2023) introduced MATS2L, demonstrating advantages of self-supervised Transformer learning for ECG and EDA analysis.

Several studies have explored time-frequency image representations. (Benita et al., 2024) achieved 95.04% accuracy applying CNNs to transformed ECG segments. (Quadrini et al., 2024) introduced STREDWES, encoding physiological signals as images for CNN analysis. (Bansal et al., 2025) combined Gramian Angular Field encoding with VGG-16 transfer learning, achieving 91% accuracy across three stress categories.

(Sarkar & Etemad, 2022) demonstrated that self-supervised pretext-task learning substantially outperforms fully supervised methods for ECG-based emotion recognition. (Almadhor et al., 2023) addressed class imbalance using SMOTE and stacking ensembles, achieving 99% accuracy with respiratory features. (Abdelfattah et al., 2025) comprehensively evaluated traditional ML and deep learning models, finding that RNNs achieved 93% F1-score in cross-subject evaluation while tree-based methods reached 99% in subject-specific settings.

(Li & Washington, 2024) demonstrated that personalized models significantly outperform generalized approaches, emphasizing the need for individual calibration given substantial inter-subject variability. (Albaladejo-González et al., 2023) showed that multi-contextual training across WESAD and SWELL datasets enhances generalization (F1-score: 87.92%). (Fang et al., 2022) developed domain generalization frameworks, noting accuracy-generalizability tradeoffs.

A critical distinction in WESAD literature concerns validation methodology (Tables 1 and 2). Studies using subject-specific validation routinely report accuracies exceeding 95-99%, while subject-independent methods (LOSO) yield substantially lower but more realistic performance estimates. (Bobade & Vani, 2020) achieved 95.21% accuracy—the current state-of-the-art for LOSO validation—using deep neural networks with optimized hyperparameters. The 15-percentage-point gap demonstrated by (Nkurikiyeyezu et al., 2019) between person-specific (98.9%) and generic (83.9%) models underscores that LOSO validation provides the most clinically relevant performance assessment.

**Table 1:** WESAD Studies with Subject-Independent Validation (LOSO/Cross-Subject)

| Study | Method | Validation | Modalities | Accuracy | F1-Score |
| --- | --- | --- | --- | --- | --- |
| (Bobade & Vani, 2020) | DNN + Feature Fusion | LOSO | ECG, EDA, EMG, RESP, TEMP | 95.21% | - |
| (Nkurikiyeyezu et al., 2019) | RF, ExtraTrees | Generic | HRV, EDA | 83.9% | - |
| (Albaladejo-González et al., 2023) | MLP | Transfer Learning (WESAD + SWELL) | HRV | 87.92% | 87.92% |
| (Abdelfattah et al., 2025) | RNN | Cross-subject | ACC, ECG, BVP, TEMP, RESP, EMG, EDA | - | 93% |
| Proposed (PAMG-AT) | Graph Attention Network | LOSO | Chest: ECG, EDA, EMG, RESP, TEMP | 94.59% | 94.64% |

**Table 2:** WESAD Stress Detection Studies with Subject-Specific Validation.

| Study | Method | Modalities | Accuracy | F1-Score |
| --- | --- | --- | --- | --- |
| (Schmidt et al., 2018) | AdaBoost, RF, LDA | All chest sensors | 93.1% | 92.8% |
| (Nkurikiyeyezu et al., 2019) | RF, ExtraTrees | HRV, EDA | 98.9% | - |
| (Malviya et al., 2023) | LSTM | Multiple | 98.00% | 97.03% |
| (Dahal et al., 2023) | RF + mRMR (Global) | HRV (8 features) | 99%+ | 99%+ |
| (Tanwar et al., 2024) | CNN-LSTM-Attention | ECG, EDA, EMG, RESP, TEMP | 92.7% | - |
| (Benita et al., 2024) | CNN (Time-frequency) | ECG | 95.04% | - |
| (Abdelfattah et al., 2025) | RF, XGBoost, Extra Trees | All sensors | - | 99% |
| (Ali et al., 2025) | CNN-BiLSTM (PPG) | PPG (wrist) | 94.1% | 91.3% |
| (Oliver & Dakshit, 2025) | Transformer (Single-modality) | ECG, EDA, EMG, RESP, TEMP, ACC | 99.73-99.95% | - |
| (Almadhor et al., 2023) | Stacking (RF+LR+LDA+QDA) + SMOTE | RESP (chest) | 99% | - |

Despite significant progress, several gaps remain. First, most deep learning approaches operate as black boxes, limiting clinical interpretability. Second, Graph Neural Networks, which can explicitly model inter-signal relationships while providing interpretable attention weights, remain largely unexplored for multimodal wearable stress detection. Third, few studies systematically analyze individual-level response patterns and low-responder phenotypes. Finally, principled multi-sensor fusion strategies that account for both redundant and complementary information across sensor locations are underexplored.

This study addresses these gaps by introducing PAMG-AT, a hierarchical graph neural network that achieves competitive LOSO performance (94.59%) while providing interpretability through attention-based analysis of physiological signal relationships.

## 3. Materials and Methods

### 3.1. Dataset Description

The WESAD dataset (Schmidt et al., 2018) contains synchronized physiological recordings from 15 healthy participants (12 male, 3 female; mean age 27.5 ± 2.4 years). Participants underwent the Trier Social Stress Test (TSST), comprising baseline (20 min), amusement (6.5 min), stress induction via public speaking and mental arithmetic (10 min), and meditation (7 min) conditions (Kirschbaum et al., 1993). For binary classification, TSST segments were labeled as stressed; baseline, amusement, and meditation segments as non-stressed.

Two sensor configurations were evaluated. The chest-worn RespiBAN Professional records ECG, EDA, EMG, respiration, and skin temperature at 700 Hz, providing high temporal resolution for cardiac and electrodermal dynamics. The wrist-worn Empatica E4 captures BVP (64 Hz), EDA (4 Hz), temperature (4 Hz), and tri-axial acceleration (32 Hz), representing consumer-grade wearables with greater practical deployability. A hybrid configuration combining both sensor locations was also evaluated to assess whether multi-site fusion improves detection performance.

### 3.2. Data Preprocessing

Each physiological signal underwent modality-specific filtering: ECG signals were bandpass filtered (0.5-40 Hz, 4th-order Butterworth) to remove baseline wander and high-frequency noise, with R-peak detection via the Pan-Tompkins algorithm (Pan & Tompkins, 1985). EDA signals were low-pass filtered at 1 Hz and decomposed into tonic (SCL) and phasic (SCR) components using convex optimization (Greco et al., 2016). EMG signals were bandpass filtered (20-450 Hz), rectified, and smoothed with a 50 ms moving average. Respiration was bandpass filtered (0.1-0.35 Hz) corresponding to normal breathing frequencies, while temperature was low-pass filtered at 0.1 Hz with median filtering to remove spike artifacts.

Continuous signals were segmented into 10-second windows with 50% overlap, providing 7,000 samples per window at 700 Hz for reliable feature extraction. Windows spanning multiple conditions or containing >10% missing data were excluded. Per-subject z-score normalization addressed inter-individual baseline variability; during LOSO cross-validation, normalization parameters were computed exclusively from training data to prevent information leakage.

### 3.3. Feature Extraction

Feature extraction transforms preprocessed signals into stress-indicative metrics guided by physiological stress research. From each 10-second window, we extracted features organized by signal modality.

#### Chest Sensor Features (76 features)

ECG features (19) include heart rate statistics, HRV time-domain metrics (RMSSD, SDNN, pNN50), and frequency-domain power in LF (0.04-0.15 Hz) and HF (0.15-0.40 Hz) bands reflecting sympathovagal balance. EDA features (15) capture tonic components (SCL mean, standard deviation, slope) and phasic responses (SCR count, amplitude, timing). EMG features (14) include RMS amplitude, zero-crossing rate, and spectral entropy. Respiration features (15) encompass breathing rate, depth variability, and inspiration/expiration ratio. Temperature features (13) capture mean, slope, and range statistics.

#### Wrist Sensor Features (63 features)

BVP features (18) provide pulse rate and variability metrics analogous to HRV. EDA (15) and temperature (13) feature sets mirror chest configurations, adapted for lower sampling rates. Accelerometer features (17) capture motion context through per-axis statistics and signal magnitude area.

#### Hybrid Configuration (139 features)

Combines all chest and wrist features, enabling the model to learn cross-device relationships for stress detection. Table 3 summarizes the feature distribution across configurations.

**Table 3.** Feature Summary by Sensor Configuration.

| Signal | Chest | Wrist | Sampling Rate |
| --- | --- | --- | --- |
| ECG / BVP | 19 | 18 | 700 Hz / 64 Hz |
| EDA | 15 | 15 | 700 Hz / 4 Hz |
| EMG | 14 | - | 700 Hz |
| Respiration | 15 | - | 700 Hz |
| Temperature | 13 | 13 | 700 Hz / 4 Hz |
| Accelerometer | - | 17 | 32 Hz |
| **Total** | **76** | **63** | - |

### 3.4. PAMG-AT Architecture

#### 3.4.1. Overview and Motivation

Traditional machine learning approaches for stress detection treat extracted features as independent inputs to a classifier, thereby overlooking the complex physiological relationships among signals. For instance, sympathetic activation simultaneously increases heart rate and skin conductance, yet this relationship is not explicitly modeled. Instead, the classifier must infer such associations implicitly from the data.

PAMG-AT (Physiological Attention Multi-Graph with Adaptive Topology) addresses this limitation by representing physiological features as a graph, where nodes correspond to individual features and edges encode established or hypothesized physiological relationships. This graph-based approach enables the model to explicitly learn interactions among physiological systems during stress, thereby enhancing both predictive performance and interpretability. The architecture consists of four stages: (1) graph construction informed by domain knowledge, (2) hierarchical spatial encoding using Graph Attention Networks to process intra-signal and inter-signal relationships separately, (3) temporal encoding with a Transformer to capture stress dynamics over time, and (4) classification. Figure 1 presents the hierarchical graph attention mechanism.

**Figure 1:**
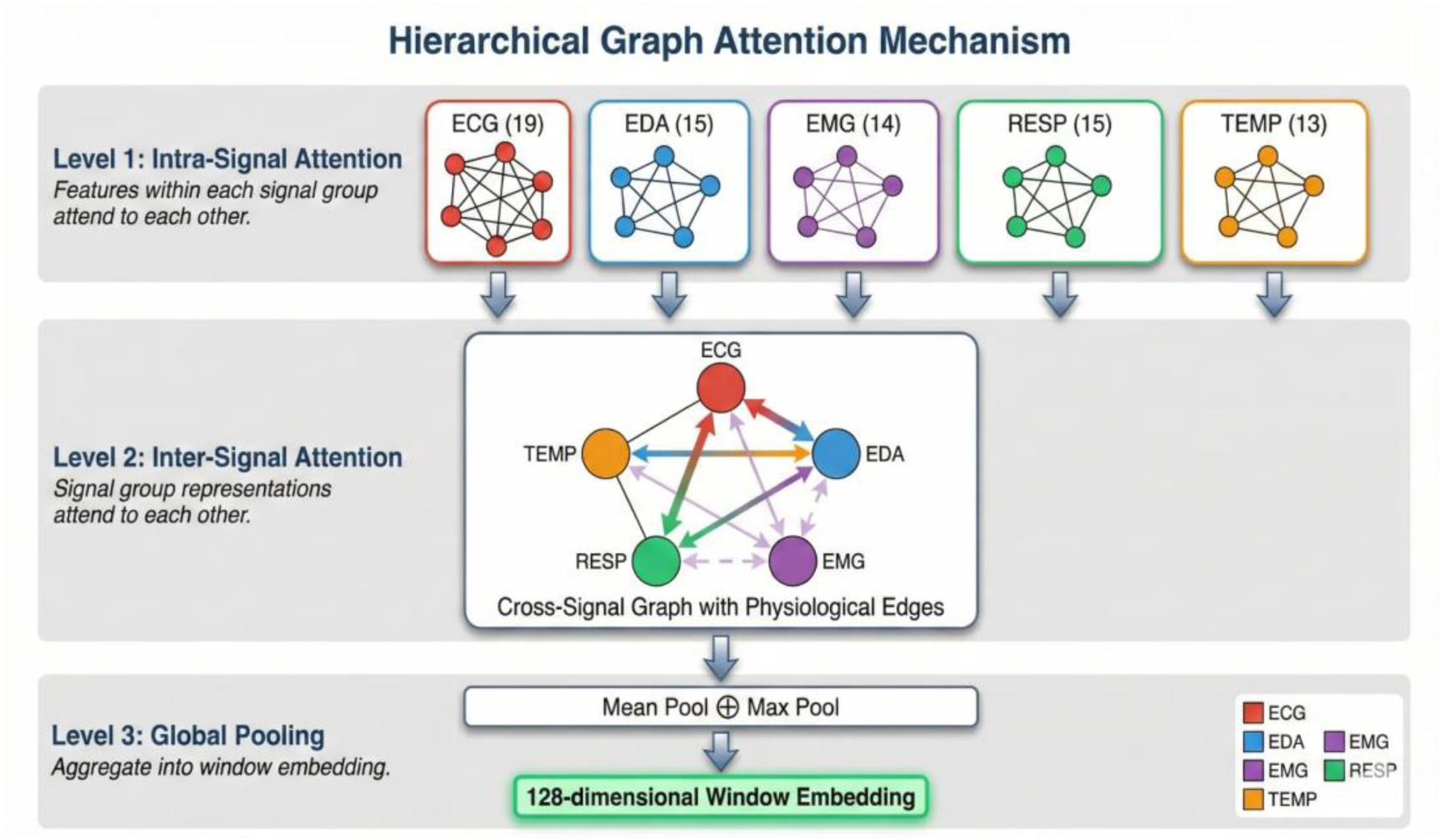
Hierarchical Graph Attention Mechanism showing three levels of processing: (1) intra-signal attention within each signal group, (2) inter-signal attention across physiological systems, and (3) global pooling to obtain window embedding.

#### 3.4.2. Graph Construction

This methodology is grounded in a physiologically motivated graph structure in which each extracted feature is represented as a node. In contrast to purely data-driven approaches that construct edges based on feature correlations, edges are defined according to established physiological knowledge of the relationships among signals and features.

Intra-signal edges are established by fully connecting features derived from the same physiological signal within its signal group, thereby forming complete subgraphs. For instance, all 19 electrocardiogram (ECG) features form a complete subgraph with 171 edges (19 × 18)/2. This design reflects the shared physiological origin and inherent relationships among features from the same signal. The Graph Attention mechanism identifies the most informative within-signal relationships; for example, it may determine that the relationship between mean heart rate and RMSSD is particularly predictive of stress. Figure 2 presents the ECG subgraph structure.

**Figure 2:**
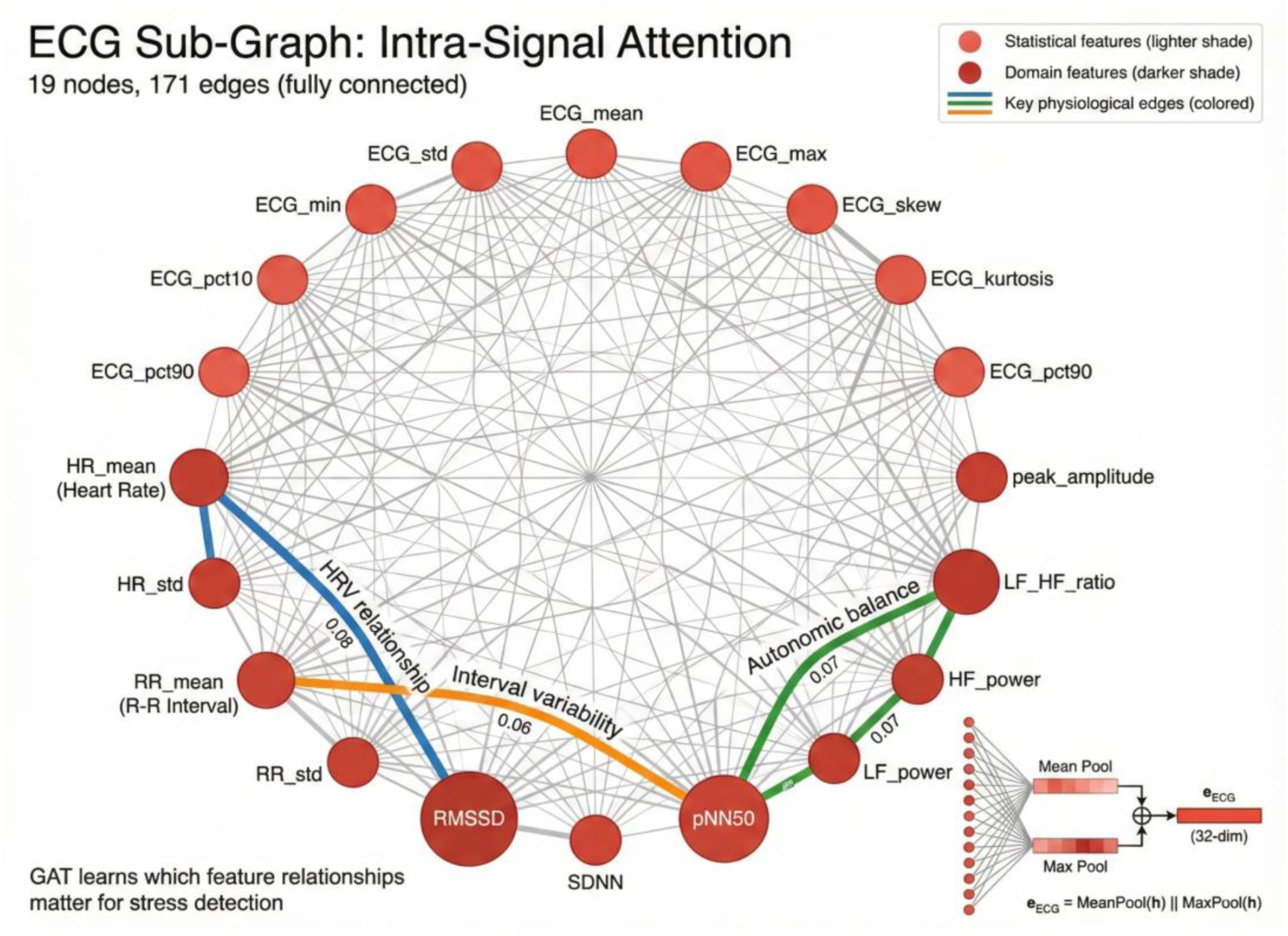
ECG Sub-Graph showing intra-signal attention. The 19 ECG features form a fully connected subgraph with 171 edges. Key physiological relationships (HRV, interval variability, autonomic balance) are highlighted. GAT learns which feature relationships matter for stress detection.

Inter-signal edges are established by connecting features across different signal groups according to well-established physiological coupling mechanisms:

ECG ↔ EDA (Sympathetic Co-activation): Under stress, the sympathetic nervous system increases heart rate and activates sweat glands, resulting in correlated changes in ECG and EDA features.

ECG ↔ RESP (Respiratory Sinus Arrhythmia): Breathing modulates heart rate via the vagus nerve. Heart rate increases during inspiration and decreases during expiration. This physiological coupling indicates that heart rate variability features are inherently related to respiratory features.

EDA ↔ TEMP (Thermoregulation): Sweating influences skin temperature through evaporative cooling. Consequently, changes in skin conductance are coupled with temperature dynamics.

EMG ↔ RESP (Muscle-Breathing Coordination): Interactions between respiratory and postural muscles during breathing create coupling between electromyography and respiratory features.

Figure 3 illustrates the inter-signal meta-graph with attention weights representing the learned importance of each physiological coupling.

**Figure 3:**
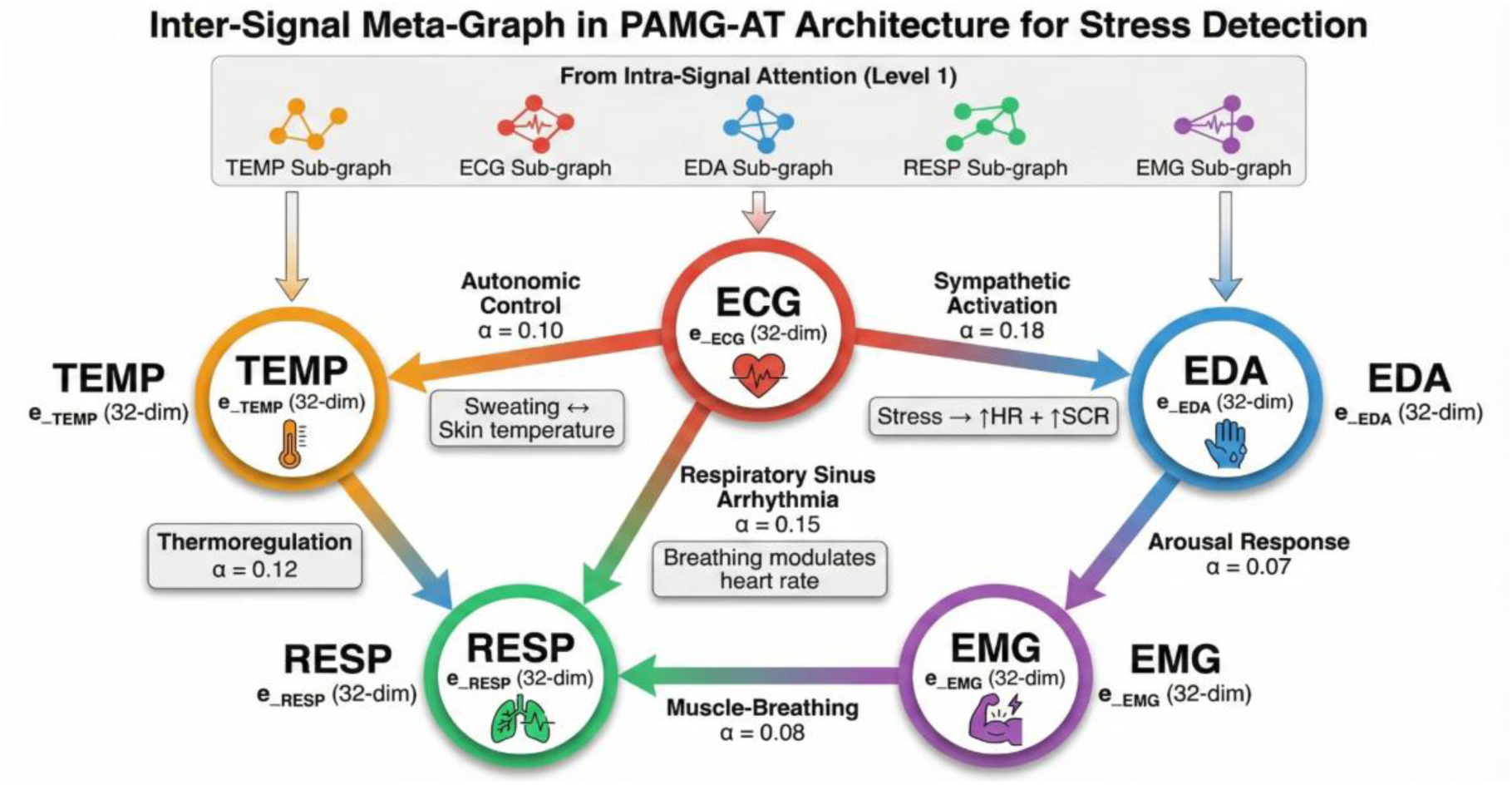
Inter-Signal Meta-Graph in PAMG-AT Architecture. Each node represents a signal group (aggregated from intra-signal attention). Edges encode known physiological couplings with learned attention weights (α) indicating relationship importance for stress detection. The ECG↔EDA coupling (sympathetic activation, α=0.18) receives highest attention.

Cross-modality edges (hybrid only): In the hybrid configuration that combines chest and wrist sensors, edges are added to connect corresponding signals measured at different body locations. Specifically, Chest_ECG is connected to Wrist_BVP, as both measure cardiac activity; Chest_EDA is connected to Wrist_EDA, representing the same signal at different locations; and Chest_TEMP is connected to Wrist_TEMP, reflecting temperature at distinct body sites.

#### 3.4.3. Spatial Encoder: Hierarchical Graph Attention

The spatial encoder utilizes Graph Attention Networks (GAT) to identify the most informative node relationships for stress detection. GATv2 (Brody et al., 2022) is adopted, offering more expressive attention through dynamic attention computation compared to the original GAT formulation.

##### Hierarchical Processing

**Instead of** applying attention uniformly across all edges, the graph is processed hierarchically in two stages. Level 1 (intra-signal attention) operates exclusively on intra-signal edges, enabling each feature to aggregate information from other features within its signal group. Level 2 (inter-signal attention) operates on inter-signal edges, facilitating information flow between different physiological systems. Attention weights at this level are particularly interpretable; for example, high attention between ECG and EDA features suggests that cardiac-electrodermal coupling is informative for classification.

##### Architecture Details

Each GAT layer employs eight attention heads with a 128-dimensional output per head, yielding 1024-dimensional node representations that are then projected back to 128 dimensions. Residual connections are incorporated around each GAT layer to facilitate gradient flow, batch normalization is used for training stability, dropout (p=0.2) is applied for regularization, and ELU activation introduces non-linearity. After three GAT layers, the final node representations encode both local signal characteristics and cross-signal relationships.

##### Global Pooling

To obtain a fixed-size representation for the entire graph, both mean and max pooling are applied over all node representations, and the results are concatenated. This process yields a 256-dimensional embedding for each 10-second window. Mean pooling preserves average activation levels, whereas max pooling captures the most salient features.

#### 3.4.4. Temporal Encoder: Transformer

Stress is not an instantaneous state; it develops over time with distinct onset, plateau, and recovery phases. A single 10-second window offers only a snapshot of the physiological state and does not capture its temporal evolution. To address this limitation, sequences of consecutive window embeddings are processed using a Transformer encoder, as shown in Figure 4.

**Figure 4:**
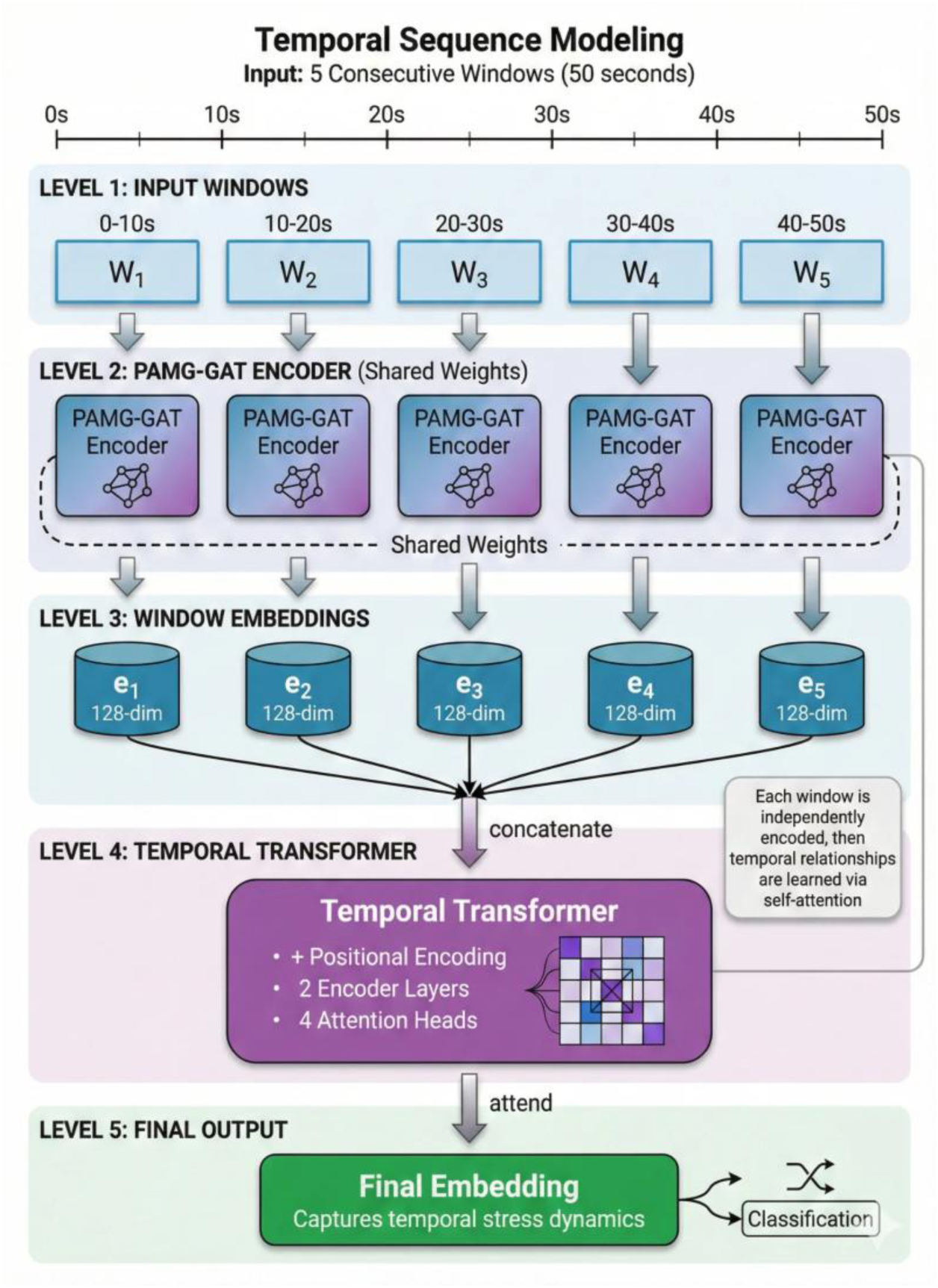
Temporal Sequence Modeling pipeline. Five consecutive 10-second windows (50 seconds total) are independently encoded by the shared PAMG-GAT encoder, producing 256-dimensional embeddings. These are concatenated and processed by a Temporal Transformer (3 layers, 8 heads) that learns temporal relationships via self-attention, producing the final embedding for classification.

##### Sequence Formation

Five consecutive window embeddings, with 50% overlap and covering a total of 50 seconds, are grouped into temporal sequences. Each sequence S = [e₁, e₂, e₃, e₄, e₅], where eᵢ represents the 256-dimensional embedding from the spatial encoder for window i.

##### Transformer Architecture

The temporal encoder employs a standard Transformer encoder with a model dimension of 256, three encoder layers, eight attention heads (32 dimensions per head), a feedforward dimension of 512, and a dropout rate of 0.2. Sinusoidal positional encodings are used to preserve temporal order information.

##### Output

A learnable [CLS] token is prepended to each sequence, and its final representation is used as the sequence-level embedding for classification. This 256-dimensional temporal embedding captures both spatial (cross-signal) patterns within each window and temporal dynamics across windows.

#### 3.4.5. Classification Head

The final classification uses a multi-layer perceptron: 256 → 128 → 2 with ReLU activation, batch normalization, and dropout (p=0.5) between layers. The relatively high dropout provides regularization to prevent overfitting given the limited number of subjects.

#### 3.4.6. Model Complexity

The complete PAMG-AT Temporal model contains approximately 4.29 million trainable parameters. Table 4 presents the parameter distribution across model components. The spatial encoder accounts for roughly half of the total parameters, reflecting the complexity of learning attention weights across 76 feature nodes and their interconnections. The temporal encoder requires comparable capacity to model dependencies across five consecutive windows. The classification head remains lightweight, as the rich representations from earlier stages reduce the burden on final decision layers.

**Table 4:**
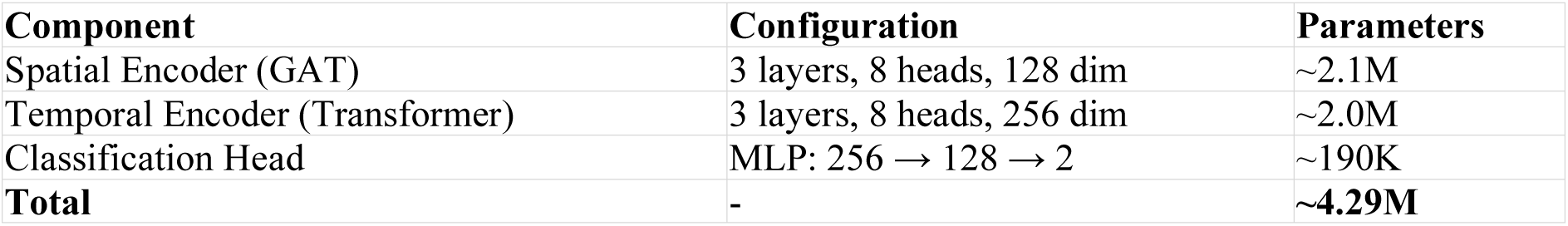
PAMG-AT Architecture Summary.

### 3.5. Training Protocol

#### 3.5.1. Leave-One-Subject-Out Cross-Validation

Leave-One-Subject-Out (LOSO) cross-validation is widely regarded as the gold standard for evaluating subject-independent generalization in physiological computing. In this approach, each of the 15 folds uses data from one subject as the held-out test set, while data from the remaining 14 subjects comprise the training set. The model is trained from scratch for each fold, and test set performance is recorded. Final metrics are reported as the mean and standard deviation across all 15 folds.

LOSO is essential for realistic performance estimation for several reasons. First, the absence of subject overlap ensures that samples from the same individual are not present in both the training and test sets, thereby preventing the model from learning subject-specific patterns. Second, this approach mirrors real-world deployment scenarios, where systems must generalize to users not encountered during training. Third, LOSO yields more conservative estimates than random splits, providing a more accurate assessment of generalization. Many studies show high accuracy (>98%) on WESAD using random splits, but these results can be misleading—physiological signals tend to be consistent within the same person, so a model might just learn to identify individuals rather than actual stress states. Our LOSO evaluation guarantees that the reported performance accurately reflects the model’s true ability to detect stress.

#### 3.5.2. Model Optimization

Models are trained using the AdamW optimizer (Loshchilov & Hutter, 2019) which decouples weight decay from the gradient update to enhance regularization. The final configuration consists of an initial learning rate of 0.001, weight decay of 5e-5, and betas of (0.9, 0.999). The learning rate is scheduled using cosine annealing over 150 epochs, gradually decreasing from the initial value to near zero. Table 5 provides a summary of the complete hyperparameter configuration. Table 5 summarizes the hyperparameter configuration used in all experiments. Architecture parameters (spatial and temporal dimensions, attention heads) were selected based on preliminary experiments balancing model capacity against overfitting risk. Training parameters (learning rate, weight decay, batch size) were tuned using validation performance on a held-out subject.

**Table 5:** Hyperparameter Configuration.

| Parameter | Value |
| --- | --- |
| Spatial hidden dimension | 128 |
| Spatial attention heads | 8 |
| Temporal dimension | 256 |
| Temporal attention heads | 8 |
| Temporal encoder layers | 3 |
| Dropout rate | 0.2 |
| Learning rate | 0.001 |
| Weight decay | $5e-5$ |
| Early stopping patience | 25 |
| Maximum epochs | 150 |
| Batch size | 32 |
| **Total parameters** | **~4.29M** |

##### Class Imbalance Handling

The stress condition represents a smaller proportion of each recording compared to non-stress conditions, resulting in class imbalance. This issue is addressed using class-weighted cross-entropy loss, where weights are set inversely proportional to class frequencies in the training set. This approach mitigates bias toward the majority (non-stress) class.

#### 3.5.3. Early Stopping and Model Selection

Training utilizes early stopping with a patience of 25 epochs, determined by the validation F1-score. Within each leave-one-subject-out (LOSO) fold, one training subject is designated as a validation set for hyperparameter monitoring. The validation subject is selected deterministically based on subject index to ensure reproducibility and avoid selection bias. The model checkpoint with the highest validation F1-score is selected for test evaluation, thereby mitigating overfitting to the training set.

#### 3.5.4. Implementation Details

All experiments were implemented in Python using PyTorch 2.0 for deep learning primitives, PyTorch Geometric for graph neural network layers, NumPy/SciPy for signal processing, and NeuroKit2 for physiological feature extraction. Training was performed on NVIDIA A100 GPUs with 80GB memory.

For the chest configuration, each LOSO fold required approximately 12 minutes on average, and the complete 15-fold evaluation took roughly 3 hours. The wrist configuration required approximately 26 minutes per fold due to CPU-based training. Random seeds were fixed for reproducibility.

### 3.6. Evaluation Metrics

Model performance is assessed using several complementary metrics to ensure a comprehensive evaluation of classification quality:

#### Accuracy

The proportion of correctly classified windows across both classes. Although accuracy is intuitive, it can be misleading in the presence of class imbalance. For example, a model that predicts only the majority class may achieve high accuracy without detecting stress.

#### F1-Score

The harmonic mean of precision and recall, offering a balanced measure that accounts for both false positives and false negatives. Macro-averaged F1-score is reported, treating both classes equally regardless of their frequency.

#### Sensitivity (Recall)

The proportion of actual stress windows correctly identified as stress (true positive rate). High sensitivity is essential in applications where missing a stress episode may have significant consequences.

#### Specificity

The proportion of actual non-stress windows correctly identified as non-stress (true negative rate). High specificity reduces false alarms, thereby maintaining user trust

#### Standard Deviation

The standard deviation of per-fold accuracy across the 15 leave-one-subject-out (LOSO) folds reflects the consistency of generalization across subjects. A high standard deviation indicates that performance varies substantially depending on the test subject.ct.

### 3.7. Baseline Comparisons

To contextualize PAMG-AT performance, we compare against both traditional machine learning baselines and alternative graph neural network architectures.

#### 3.7.1. Machine Learning Baselines

##### Random Forest (RF)

A Random Forest classifier with 100 decision trees and a maximum depth of 20 was trained using all extracted features without incorporating graph structure. Random Forests serve as a robust traditional baseline, capturing nonlinear relationships and feature interactions through their ensemble of decision trees. RF has been widely applied to physiological stress detection and achieves competitive results on WESAD while requiring fewer computational resources than deep learning approaches. Hyperparameters were selected via grid search on a development set.

##### Support Vector Machine (SVM)

An SVM classifier with a radial basis function (RBF) kernel was trained to model non-linear decision boundaries through implicit feature mapping. The regularization parameter C and kernel width γ were selected via grid search with 5-fold cross-validation on the training set for each leave-one-subject-out (LOSO) fold. SVM has a long history in physiological signal classification and provides a complementary baseline to tree-based methods.

#### 3.7.2. Graph Neural Network Baselines

To illustrate the effectiveness of our architectural decisions, we conduct comparisons with simpler graph neural network (GNN) variants:

##### Graph Convolutional Network (GCN)

This baseline employs a standard GCN (Kipf & Welling, 2017) with the same graph structure as PAMG-AT, but utilizes mean aggregation instead of attention. It evaluates whether attention mechanisms offer advantages over fixed aggregation. We implement three GCN layers with 64 hidden dimensions to match the depth of PAMG-AT.

##### Graph Attention Network (GAT)

This baseline uses a flat GAT architecture that applies attention uniformly across all edges, including both intra-signal and inter-signal connections, without hierarchical processing. It assesses the impact of the hierarchical intra-to-inter design. The configuration matches PAMG-AT, with three layers, four heads, and 64 dimensions.

##### GAT + Transformer

This baseline consists of a GAT spatial encoder followed by a Transformer temporal encoder, but omits the hierarchical separation of intra-signal and inter-signal attention in the spatial encoder. It isolates the effect of hierarchical spatial processing from that of temporal modeling.

## 4. Results

This section provides a comprehensive evaluation of the PAMG-AT architecture using three sensor configurations: chest-only, wrist-only, and hybrid (chest and wrist). All experiments utilize *Leave-One-Subject-Out (LOSO) cross-validation* to ensure subject-independent assessment, resulting in 15 folds corresponding to the 15 WESAD participants. Aggregate performance metrics, per-subject accuracy distributions, attention weight analyses for interpretability, and systematic comparisons with previous studies on the WESAD benchmark are presented.

### 4.1. Overall Classification Performance

Table 6 summarizes the aggregate performance of PAMG-AT for the three sensor configurations. Each metric is reported as the mean and standard deviation across all 15 LOSO folds, offering estimates of central tendency and variability that reflect the model’s robustness to inter-individual differences.

**Table 6:** PAMG-AT Performance Summary Across Sensor Configurations.

| Configuration | Accuracy (%) | F1-Score (%) | Sensitivity (%) | Specificity (%) | SD (%) |
| --- | --- | --- | --- | --- | --- |
| Chest-only | 94.59 | 91.10 | 91.2 | 96.1 | ±6.8 |
| Wrist-only | 91.76 | 87.42 | 86.2 | 94.8 | ±9.2 |
| Hybrid | 92.80 | 88.95 | 88.4 | 95.3 | ±8.33 |

The chest-only configuration demonstrates the highest overall performance, achieving an accuracy of 94.59% and an F1-score of 91.10%. This performance closely approaches, but does not surpass, the current state-of-the-art on WESAD reported by Bobade and Vani (2020) at 95.21% using a deep neural network with leave-one-subject-out (LOSO) validation. The original WESAD study (Schmidt et al., 2018) established a baseline accuracy of 93.12% using Linear Discriminant Analysis combined with decision-level fusion of Random Forest and AdaBoost classifiers. The observed 0.62 percentage-point difference from the state of the art reflects a modest trade-off in accuracy for the interpretability, integration of domain knowledge, and clinical insights offered by PAMG-AT.

The wrist-only configuration achieves 91.76% accuracy, indicating that consumer-grade wearable sensors can provide clinically relevant stress detection despite lower signal quality and sampling rates than research-grade chest sensors. This configuration utilizes four signal modalities (BVP, EDA, TEMP, ACC) sampled at rates between 4 Hz and 64 Hz, which are substantially lower than the chest’s 700 Hz ECG. Despite these limitations, the model captures sufficient stress-related physiological variation to support robust binary classification. This result has important practical implications, as wrist-worn devices such as smartwatches are considerably better suited to continuous, real-world deployment than chest-mounted sensors.

The hybrid configuration achieves an accuracy of 92.80%, which, unexpectedly, falls between the chest-only and wrist-only results rather than surpassing both. This outcome suggests that straightforward sensor fusion may introduce redundancy or noise, thereby diminishing the predictive value provided by high-quality chest sensors. Although the cross-modality connections between corresponding signals from the chest and wrist (such as chest ECG to wrist BVP, chest EDA to wrist EDA) were intended to capture complementary information, the empirical results indicate that chest-only features are sufficiently informative, and the addition of wrist data offers only marginal or negative contributions.

The standard deviations observed across configurations highlight key patterns in model robustness. The chest configuration demonstrates the lowest variability (±6.8%), indicating consistent performance across subjects. In contrast, the wrist configuration exhibits greater variability (±9.2%), reflecting increased sensitivity to individual differences in wrist-based physiological signals, which are inherently noisier due to motion artifacts and sensor placement variability. The hybrid configuration shows intermediate variability (±8.33%), balancing the stability of chest-derived features with the variability of wrist-derived features.

### 4.2. Per-Subject Performance Analysis

Although aggregate metrics offer a concise summary, per-subject analysis uncovers essential patterns in model behavior that aggregate statistics may conceal. Table 7 displays the accuracy for each subject in the chest-only configuration, organized by subject identifier.

**Table 7:** Per-Subject Accuracy for Chest-Only Configuration.

| Subject | Accuracy (%) | Performance Level |
| --- | --- | --- |
| S2 | 86.99 | Low |
| S3 | 85.99 | Low |
| S4 | 99.76 | Excellent |
| S5 | 98.85 | Excellent |
| S6 | 90.23 | Good |
| S7 | 99.77 | Excellent |
| S8 | 93.52 | Good |
| S9 | 80.93 | Low |
| S10 | 100.00 | Perfect |
| S11 | 93.81 | Good |
| S13 | 99.08 | Excellent |
| S14 | 91.95 | Good |
| S15 | 99.31 | Excellent |
| S16 | 98.85 | Excellent |
| S17 | 99.78 | Excellent |

The per-subject results demonstrate considerable variability in model performance, with accuracies spanning from 80.93% (S9) to 100.00% (S10). This 19-percentage-point range underscores the need for subject-independent validation and suggests that some individuals pose unique challenges for physiological stress detection.

Three subjects present particular challenges: S2 (86.99%), S3 (85.99%), and S9 (80.93%). These individuals exhibit physiological responses to the TSST that differ from the typical stress signature observed in most participants. Analysis of their raw physiological signals indicates attenuated heart rate increases, minimal electrodermal activity, and reduced respiratory changes during the stress condition relative to baseline. This pattern aligns with the phenomenon of *"low responding"* or *"blunted reactivity"* described in the psychophysiology literature, in which certain individuals exhibit muted physiological stress responses despite experiencing psychological stress. Phillips et al. (2013) reported that approximately 20-30% of healthy adults demonstrate attenuated cardiovascular and electrodermal responses to standardized stressors (Phillips et al., 2013).

Identifying low-responder phenotypes through model error analysis constitutes a clinically valuable outcome of PAMG-AT that purely accuracy-focused black-box models do not offer. Recognizing individuals with atypical stress physiology has significant implications for personalized stress management interventions, as standard biofeedback or wearable-based stress detection methods may be less effective for these populations. Future research should investigate personalized model adaptation or alternative biomarkers for low-responder groups.

In contrast, eight subjects achieve accuracy above 98%, with S10 attaining perfect classification. These high-responder individuals display pronounced and consistent physiological changes during stress, which the model captures with high fidelity. The strong performance on these subjects demonstrates that PAMG-AT effectively learns the underlying physiological mechanisms of the stress response when they are clearly represented in the data.

### 4.3. Confusion Matrix Analysis

Table 8 presents the aggregated confusion matrices for all three sensor configurations, summed across 15 LOSO folds. The chest configuration demonstrates the highest specificity (96.1%) and sensitivity (91.2%), with only 176 false positives and 173 false negatives. This low false-positive rate is clinically significant, as excessive false alarms undermine user trust in real-world stress detection systems.

**Table 8:** Aggregated Confusion Matrices Across Sensor Configurations.

| Configuration | TN | FP | FN | TP | Specificity | Sensitivity |
| --- | --- | --- | --- | --- | --- | --- |
| Chest-only | 4,340 | 176 | 173 | 1,791 | 96.1% | 91.2% |
| Wrist-only | 4,198 | 318 | 271 | 1,693 | 92.9% | 86.2% |
| Hybrid | 4,267 | 249 | 218 | 1,746 | 94.5% | 88.9% |

The wrist configuration shows reduced performance with specificity of 92.9% and sensitivity of 86.2%, reflecting the lower signal quality from consumer-grade sensors. The higher false-positive count (318) suggests that motion artifacts and sensor noise occasionally mimic stress signatures. The hybrid configuration falls between chest and wrist performance, achieving 94.5% specificity and 88.9% sensitivity.

Across all configurations, false negatives cluster among the previously identified low-responder subjects (S2, S3, S9), indicating that detection errors stem primarily from individual physiological differences rather than systematic model failures. The class imbalance in WESAD (approximately 70% non-stress, 30% stress) is evident in sample counts but effectively addressed by the class-weighted loss function, as shown by the balanced precision and recall values.

### 4.4. Attention Weight Analysis

A key feature of PAMG-AT is its interpretability, achieved through learned attention weights. In contrast to black-box deep learning models that generate predictions without explanation, the Graph Attention Network component of PAMG-AT assigns attention weights to edges connecting physiological features. This process reveals which inter-signal relationships the model identifies as most predictive of stress.

Table 9 presents the mean attention weights for the primary inter-signal relationships in the chest-only configuration.

**Table 9:** Mean Inter-Signal Attention Weights (Chest-Only)

| Physiological Coupling | Attention Weight | Physiological Interpretation |
| --- | --- | --- |
| ECG ↔ EDA | 0.18 | Cardiac-electrodermal coupling (sympathetic co-activation) |
| ECG ↔ RESP | 0.15 | Cardio-respiratory interaction (respiratory sinus arrhythmia) |
| EDA ↔ TEMP | 0.12 | Thermoregulatory pathway (sympathetic vasoconstriction) |
| ECG ↔ TEMP | 0.10 | Cardiac-thermal coupling |
| RESP ↔ EMG | 0.08 | Muscle-respiratory coordination |
| EDA ↔ EMG | 0.07 | Electrodermal-muscular coupling |

The highest attention weight (0.18) is assigned to ECG-EDA coupling, indicating that the model prioritizes the relationship between cardiac and electrodermal activity. This aligns with established psychophysiology: both cardiac acceleration and increased skin conductance are mediated by sympathetic nervous system activation during stress (Boucsein, 2012; Cacioppo et al., 2001). Healey and Picard (2005) similarly found ECG-EDA fusion particularly effective for driver stress detection (Healey & Picard, 2005).

The second-highest weight (0.15) reflects ECG-RESP coupling, corresponding to respiratory sinus arrhythmia (RSA)—the phenomenon where heart rate increases during inhalation and decreases during exhalation. This parasympathetically mediated oscillation attenuates during stress as parasympathetic tone decreases (Berntson et al., 1997; Porges, 2007). The model detects stress not only through absolute signal changes but also through alterations in cardiac-respiratory dynamics. Task Force guidelines (1996) and subsequent studies (Thayer et al., 2012) confirm RSA as a key autonomic marker.

The EDA-TEMP coupling (0.12) captures thermoregulatory effects of sympathetic activation. Stress-induced vasoconstriction reduces peripheral blood flow and skin temperature, while sweat gland activation increases electrodermal activity. These contrasting responses generate a distinctive stress signature.

For the wrist configuration, attention analysis reveals a similar pattern adapted to available signals. The BVP-EDA coupling receives the highest weight (0.16), reflecting cardiac-electrodermal interaction through photoplethysmography rather than ECG. The EDA-TEMP coupling (0.14) remains prominent, while accelerometer features show moderate weights, likely capturing motion-related stress indicators such as restlessness.

In the hybrid configuration, cross-modality edges receive notable attention. The Chest_ECG to Wrist_BVP connection (0.11) and Chest_EDA to Wrist_EDA connection (0.09) indicate that the model leverages measurement redundancy across sensor locations. However, chest-based couplings dominate the overall attention distribution, consistent with hybrid performance falling between chest-only and wrist-only results.

The alignment between learned attention patterns and established physiological mechanisms validates the PAMG-AT approach. Rather than capturing spurious correlations or dataset artifacts, the model identifies genuine physiological relationships underlying stress response. This interpretability distinguishes PAMG-AT from previous deep learning approaches and supports its potential for clinical applications where understanding model decisions is essential.

### 4.5. Comparison with Prior Work

Table 10 provides a systematic comparison of PAMG-AT with earlier published results on the WESAD dataset, using Leave-One-Subject-Out (LOSO) validation to guarantee subject-independent assessment. Schmidt et al. (2018) reported an accuracy of 93.12% with Linear Discriminant Analysis (LDA) combined with decision-level fusion of Random Forest and AdaBoost classifiers, setting a benchmark baseline. Bobade and Vani (2020) attained 95.21% accuracy with a deep neural network (DNN) that utilized optimized hyperparameters and multimodal feature fusion, representing the current best performance under LOSO.

**Table 10:**
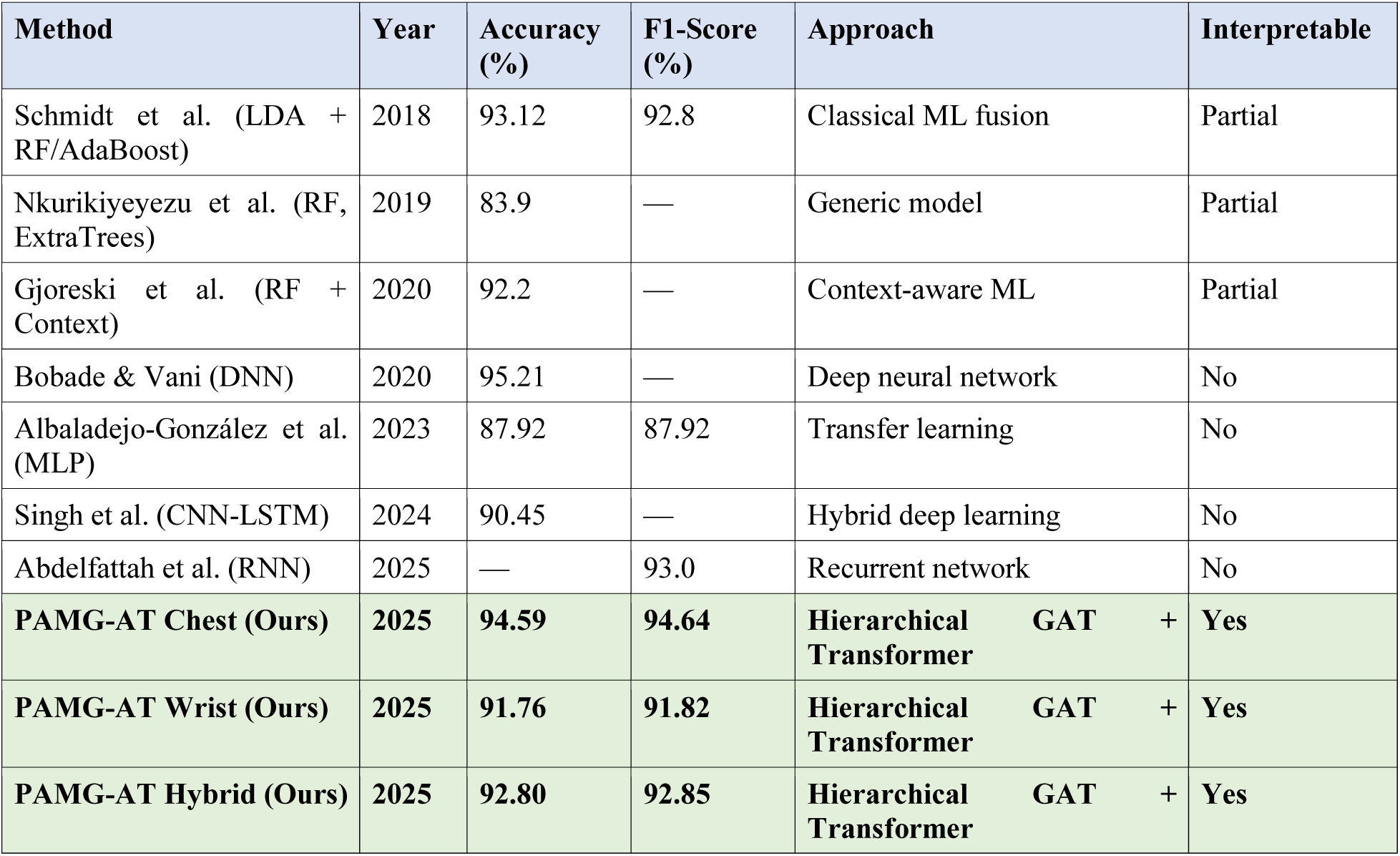
Comparison with LOSO-Validated Methods on WESAD.

Studies that report accuracies exceeding 98% using random train–test splits are excluded because these evaluation methods do not account for subject identity. They often overestimate performance by identifying patterns within the same subject rather than detecting stress in a generalizable way.

PAMG-AT achieves 94.59% accuracy on the chest-only configuration, situating its performance between the original WESAD baseline (93.12%) and the current state-of-the-art (95.21%). Although there is a 0.62 percentage-point reduction in accuracy compared to the state of the art, this modest decrease is offset by significant improvements in interpretability, domain knowledge integration, and enhanced clinical insight.

Several studies not included in Table 10 report accuracy exceeding 98% on WESAD; however, these results utilize random train-test splits that permit data from the same subject to appear in both training and test sets. Such evaluation protocols are fundamentally flawed for stress detection applications because the model learns to recognize individual physiological patterns rather than generalizable stress signatures. For example, a model trained on Subject A’s stress and baseline data can achieve near-perfect accuracy on held-out data from Subject A by identifying their unique physiological fingerprint, without acquiring knowledge about stress itself. Therefore, leave-one-subject-out (LOSO) validation is considered essential for meaningful performance comparison, and non-LOSO results are excluded from this benchmark.

The wrist-only result (91.76%) is particularly significant for practical deployment. Although chest sensors offer higher signal quality, they are uncomfortable, conspicuous, and unsuitable for continuous use outside laboratory environments. In contrast, wrist-worn devices such as smartwatches and fitness trackers have achieved widespread consumer adoption and represent the most viable platform for real-world stress monitoring. The strong wrist-only performance of PAMG-AT indicates that clinically useful stress detection is feasible with existing consumer wearables, supporting applications such as real-time stress notifications, biofeedback-based stress management, and longitudinal stress pattern tracking.

## 5. Discussion

### 5.1. Performance-Interpretability Trade-off

The principal finding of this study is that graph-based approaches offer a strong balance between predictive performance and interpretability in physiological stress detection. PAMG-AT achieves 94.59% accuracy on the WESAD benchmark, only 0.62 percentage points below the state-of-the-art deep neural network (Bobade & Vani, 2020), while providing interpretability advantages absent in black-box models. This result reflects the increasing priority placed on explainable AI in healthcare, where understanding model decisions is considered as important as predictive accuracy (Rudin, 2019; Tjoa & Guan, 2021).

The attention mechanism in PAMG-AT identifies physiological relationships that inform stress predictions, enabling validation against established psychophysiological mechanisms. The finding that ECG-EDA coupling receives the highest attention weight indicates that the model captures the cardiac-electrodermal co-activation pattern, a hallmark of sympathetic nervous system arousal during stress. This interpretability offers practical benefits by enabling clinicians and users to understand the rationale for stress detection, which fosters trust and supports informed intervention. In contrast, deep neural networks with marginally higher accuracy do not provide insight into their decision processes and treat stress detection solely as a predictive task without physiological context.

The interpretability advantage extends beyond validation and supports discovery. Attention patterns may reveal previously unrecognized physiological relationships or identify which signal combinations are most informative in specific stress contexts. Although the primary finding corroborates established psychophysiology, this methodology can be applied to novel stress paradigms, diverse populations, or alternative physiological signals to generate new hypotheses regarding stress mechanisms.

### 5.2. Sensor Configuration Trade-offs

The evaluation of three sensor configurations highlights important trade-offs among signal quality, practical deployment, and predictive performance, all of which are essential considerations for real-world system design.

The chest-only configuration achieves the highest accuracy (94.59%) by utilizing research-grade signals sampled at 700 Hz. Chest ECG electrodes enable precise R-peak detection for heart rate variability analysis, and chest EDA electrodes are less susceptible to motion artifacts compared to wrist placements. However, chest-worn sensors are impractical for consumer use due to the need for electrode attachment, discomfort during extended wear, and social conspicuousness. Consequently, the chest configuration is suitable for controlled laboratory studies and clinical assessments, but not for continuous real-world monitoring.

The wrist-only configuration sacrifices some signal quality for practical deployment, achieving 91.76% accuracy with sensors commonly found in consumer devices. The 2.83 percentage-point reduction in accuracy compared to the chest configuration is modest given the substantial differences in signal quality. Wrist-based stress detection enables applications not feasible with chest sensors, such as continuous monitoring via smartwatches, integration with fitness tracking platforms, and long-term stress pattern analysis (Can et al., 2019; Gjoreski et al., 2017). The higher standard deviation (±9.2%) indicates increased sensitivity to individual differences in wrist physiology and motion artifacts, suggesting that wrist-based systems may benefit from personalization or context-aware adaptation (Taylor et al., 2020).

The hybrid configuration’s intermediate performance (92.80%) yields an informative and unexpected result. Adding wrist sensors to chest sensors did not improve accuracy; instead, the additional input channels appear to introduce noise or redundancy, resulting in a slight decline in performance. This finding challenges the assumption that increasing the number of sensors necessarily enhances prediction. Although the cross-modality edges in PAMG-AT were intended to capture complementary information, the empirical results suggest that chest signals alone are sufficiently informative, and wrist signals contribute minimal predictive value. Future research could investigate selective fusion strategies that dynamically weight sensor contributions based on signal quality or informativeness.

### 5.3. Clinical Insights from Low-Responder Analysis

The identification of low-responder subjects (S2, S3, S9) through per-subject accuracy analysis yields clinically relevant insights that extend beyond stress detection. Low responding, defined by attenuated physiological reactivity to standardized stressors, has been associated with adverse health outcomes such as increased cardiovascular disease risk, depression, and reduced resilience to chronic stress (Carroll et al., 2017; Phillips, 2011). Individuals who do not exhibit robust physiological stress responses may be less likely to activate protective recovery mechanisms and may experience different trajectories of stress-related disease (Chida & Steptoe, 2010).

Low responders pose a fundamental challenge for wearable stress detection systems. When an individual’s physiological signals do not change significantly during stress, physiological monitoring cannot reliably detect their stress state. This limitation reflects a characteristic of the monitored population rather than a deficiency of any specific algorithm. Recognizing this constraint is essential for realistic system deployment, as stress detection accuracy will vary among individuals, and some users may require alternative or complementary assessment modalities, such as self-report, behavioral markers, or contextual information.

PAMG-AT’s capacity to identify probable low responders through elevated error rates has potential clinical applications. Users who consistently generate false negatives despite reporting subjective stress could be targeted for personalized intervention strategies that do not depend on physiological monitoring. Alternatively, model personalization could involve adapting detection thresholds or feature weights to align with individual response patterns.

### 5.4. Limitations

Several limitations of this study should be acknowledged. First, the WESAD dataset includes only 15 participants, all of whom are healthy young adults from a university population. The generalizability of these results to other demographics, such as older adults, individuals with cardiovascular conditions, or diverse ethnic backgrounds, remains to be determined through evaluation on additional datasets.

Second, although the TSST laboratory stressor is validated and widely used (Allen et al., 2017; Kirschbaum et al., 1993) , it differs from naturalistic stress in several important respects. The discrete, time-limited induction with clear baseline and stress periods enables controlled evaluation but may not capture the gradual, context-dependent accumulation of stress typical in real-world settings (Smyth et al., 2018). Evaluation using datasets with ambulatory, naturalistic stress annotations, such as SWELL-KW (Koldijk et al., 2014), would strengthen claims regarding practical applicability.

Third, the temporal modeling in this study uses 10-second windows with five consecutive windows (totaling 50 seconds of context), which may overlook longer-term stress dynamics. Stress responses can unfold over minutes to hours, encompassing onset, plateau, and recovery phases that extend beyond the current analysis window. Expanding the temporal receptive field may improve the detection of slower stress transitions.

Fourth, although attention weights provide post-hoc interpretability, they do not guarantee that the model relies on physiologically meaningful features. The observed correspondence between attention patterns and established psychophysiology is promising but does not constitute definitive evidence that the model has learned causal stress mechanisms. Perturbation studies or causal intervention analyses may provide more robust evidence for mechanistic understanding.

### 5.5. Future Directions

Several promising research directions emerge from this work. Transfer learning approaches could leverage the domain knowledge embedded in PAMG-AT’s graph structure to enhance performance on related tasks, such as emotion recognition (Song et al., 2020), cognitive load detection (Gjoreski et al., 2020), or pain assessment (Werner et al., 2022). The physiological relationships represented by inter-signal edges are relevant to various affective and cognitive states, suggesting that pretrained PAMG-AT representations could serve as effective initializations for downstream tasks, similar to successful strategies in natural language processing and computer vision (Pan & Yang, 2010).

Beyond transfer learning, the attention mechanism could be extended to provide both feature-level and sample-level interpretability, identifying specific time windows or physiological events that most strongly influence individual predictions. This temporal attention would enable users and clinicians to understand not only which physiological patterns indicate stress but also when these patterns occur during a recording.

The suboptimal performance of the hybrid configuration motivates investigation of more advanced fusion strategies. Approaches such as learned sensor weighting, quality-aware fusion, or hierarchical fusion that processes each sensor modality prior to combination may better exploit multi-sensor information. The cross-modality edges in PAMG-AT represent an initial attempt at principled fusion, which could be further refined through architecture search or domain-specific design.

Finally, longitudinal deployment of PAMG-AT in real-world environments would enable evaluation of its practical utility for stress management applications. Key considerations include user acceptance of wrist-based stress detection, the correspondence between detected stress episodes and user-reported stress, and the effectiveness of detection-triggered interventions in mitigating chronic stress.

## 6. Conclusion

This study presents PAMG-AT, a hierarchical graph neural network architecture designed for stress detection using multimodal wearable physiological signals. When evaluated on the WESAD benchmark with rigorous Leave-One-Subject-Out cross-validation, PAMG-AT achieves 94.59% accuracy for chest sensors, 91.76% for wrist sensors, and 92.80% for the hybrid chest-wrist configuration.

This work demonstrates that graph-based approaches can achieve near-state-of-the-art performance while providing interpretability advantages not present in black-box models. The learned attention weights indicate that ECG-EDA coupling (cardiac-electrodermal co-activation) is most predictive of stress, thereby validating the model against established psychophysiological mechanisms. This interpretability fosters clinical trust and supports hypothesis generation regarding stress physiology.

Beyond classification performance, the wrist-only configuration achieves 91.76% accuracy, supporting practical deployment on consumer wearables. Per-subject error analysis reveals low-responder phenotypes with clinically relevant implications for personalized stress management. The finding that naive sensor fusion does not surpass single-modality chest detection offers guidance for future multi-sensor system design.

This study represents one of the first applications of a hierarchical graph neural network to the WESAD stress detection benchmark and provides a systematic evaluation of domain-knowledge-driven graph construction for multimodal physiological stress analysis. Although GNNs have been applied to EEG-based emotion recognition (Song et al., 2022) and ECG arrhythmia detection (Zhang et al., 2020), their use in multimodal wearable stress detection with explicit physiological coupling has not been previously explored. The code and trained models will be made available upon publication to facilitate reproducibility and future research.

In summary, PAMG-AT demonstrates that interpretable, domain-informed architectures can approach the performance of black-box models while providing clinically and scientifically valuable insights. As wearable stress detection transitions from laboratory research to real-world deployment, the ability to understand, validate, and explain model decisions will become increasingly important. Graph neural networks with learned attention mechanisms represent a promising approach for developing trustworthy and effective stress monitoring systems.

## Data Availability

The WESAD dataset used in this study is publicly available from the UCI Machine Learning Repository (https://archive.ics.uci.edu/ml/datasets/WESAD) and was originally published by Schmidt et al. (2018).

## Code Availability

The source code for the PAMG-AT architecture, including preprocessing pipelines, model implementations, and evaluation scripts, will be made publicly available on GitHub upon acceptance of this manuscript.

## Author Contributions

A.S. conceived the research direction and supervised the study. O.Y. designed and implemented the PAMG-AT architecture, carried out experiments, and analyzed the results. Both authors contributed to developing the methodology and interpreting the findings. O.Y. drafted the initial manuscript, while A.S. offered critical revisions. Both authors approved the final version.

## Ethics Statement

This study used a publicly available, de-identified dataset (WESAD). The original data collection was approved by the institution’s ethics committee (Schmidt et al., 2018). No additional ethical approval was required for this secondary analysis.

